# Role of VapBC4 toxin-antitoxin system of *Sulfolobus acidocaldarius* in heat stress adaptation

**DOI:** 10.1101/2024.06.06.597757

**Authors:** Arghya Bhowmick, Alejandra Recalde, Chandrima Bhattacharyya, Jagriti Das, Ulises E. Rodriguez-Cruz, Sonja-Verena Albers, Abhrajyoti Ghosh

## Abstract

Toxin-antitoxin (TA) systems are important for stress adaptation in prokaryotes, including persistence, antibiotic resistance, pathogenicity, and biofilm formation. Toxins can cause cell death, reversible growth stasis, and direct inhibition of crucial cellular processes through various mechanisms, while antitoxins neutralize the effects of toxins. In bacteria, these systems have been studied in detail, whereas their function in archaea remains elusive. During heat stress, the thermoacidophilic archaeon *Sulfolobus acidocaldarius* exhibited an increase in the expression of several bicistronic type II *vapBC* TA systems, with the highest expression observed in the *vapBC4* system. In the current study, we performed a comprehensive biochemical characterization of the VapBC4 TA system, establishing it as a bonafide type II toxin-antitoxin system. The VapC4 toxin is shown to have high-temperature catalyzed RNase activity specific for mRNA and rRNA, while the VapB4 antitoxin inhibits the toxic activity of VapC4 by interacting with it. VapC4 toxin expression led to heat-induced persister-like cell formation, allowing the cell to cope with the stress. Furthermore, this study explored the impact of *vapBC4* deletion on biofilm formation, whereby deletion of *vapC4* led to increased biofilm formation, suggesting its role in regulating biofilm formation. Thus, during heat stress, the liberated VapC4 toxin in cells could potentially signal a preference for persister cell formation over biofilm growth. Thus, our findings shed light on the diverse roles of the VapC4 toxin in inhibiting translation, inducing persister cell formation, and regulating biofilm formation in *S. acidocaldarius*, enhancing our understanding of TA systems in archaea.

**IMPORTANCE:** This research enhances our knowledge of Toxin-antitoxin (TA) systems in archaea, specifically in the thermoacidophilic archaeon *Sulfolobus acidocaldarius*. TA systems are widespread in both bacterial and archaeal genomes, indicating their evolutionary importance. However, their exact functions in archaeal cellular physiology are still not well understood. This study sheds light on the complex roles of TA systems and their critical involvement in archaeal stress adaptation, including persistence and biofilm formation. By focusing on *S. acidocaldarius*, which lives in habitats with fluctuating temperatures that can reach up to 90℃, the study reveals the unique challenges and survival mechanisms of this organism. The detailed biochemical analysis of the VapBC4 TA system, and its crucial role during heat stress, provides insights into how extremophiles can survive in harsh conditions. The findings of this study show the various functions of the VapC4 toxin, including inhibiting translation, inducing persister-like cell formation, and regulating biofilm formation. This knowledge improves our understanding of TA systems in thermoacidophiles and has broader implications for understanding how microorganisms adapt to extreme environments.

## INTRODUCTION

In recent years, the investigation of toxin-antitoxin (TA) systems has gained attention due to their essential roles in regulating bacterial growth, aiding survival under stressful conditions, and their potential implications in bacterial virulence and antibiotic resistance [1, 2]. TA systems consist of a pair of closely linked genes encoding a stable toxin and a labile antitoxin, wherein the antitoxin neutralizes the toxin’s effects, maintaining cellular equilibrium [3]. While extensively studied in bacteria, the presence and functional significance of these systems in archaea remain relatively unexplored. Virulence-associated proteins (Vaps) constitute a distinctive subset within the type-II toxin-antitoxin (TA) systems. They exhibit a genetic arrangement with two linked genes, where a proteolytically unstable antitoxin (VapB) is commonly encoded first, followed by a stable ribonucleolytic toxin (VapC). Initially associated with virulence plasmids in human pathogens, like *Salmonella dublin*, these proteins have gained recognition as a prominent variant of type-II toxins, particularly prevalent among archaea [4, 5]. The significance of the presence of a large number of type II vapBC TA systems in archaea, which have not shown any signs of virulence, is yet to be determined. The functional scope of VapC toxins in bacteria encompasses the degradation of diverse RNA molecules like mRNA, rRNA, and tRNA [6–11]. However, interestingly, tRNAs have emerged as the primary RNA targets for most VapCs [12].

*Sulfolobus acidocaldarius*, a thermoacidophilic archaeon, thrives in a challenging environment of hot mud pools and solfataric springs where temperatures range from 75℃ to 80℃ and the pH levels are highly acidic (pH 2-3) [13]. The temperature of these habitats is constantly fluctuating owing to the geothermal activities of the earth’s crust, with the potential to reach as high as 90℃, presenting a serious threat to the resident organisms. Surviving under such extreme conditions requires utilizing intricate molecular strategies, rendering *S. acidocaldarius* a model organism for investigating unique survival mechanisms [14]. Recent genomic analyses have identified multiple TA systems in the *S. acidocaldarius* genome, with the majority being type II TA systems [15, 16]. These type II TA systems comprise a protein toxin targeting specific cellular processes, counteracted by a corresponding antitoxin protein. Furthermore, within the phylum of thermoacidophilic thermoproteota the *vapBC* system stands out as the most encountered type II TA system [16]. Previous studies by Cooper et al. demonstrated that with the increase in the optimal growth temperature of archaea, there is a concurrent increase in the count of *vapBC* TA genes within the genome of archaea [17]. This correlation implies that the *vapBC* TA system could enable archaea to adapt to higher temperatures. Also, disruption of the *vapBC*6 operon from *S. solfataricus* through targeted gene inactivation resulted in two recessive phenotypes associated with fitness: heightened sensitivity to high temperatures and a heat-dependent decrease in the growth rate [18].

The VapBC TA systems play critical roles in regulating cellular processes, stress responses, and survival strategies. In bacteria, VapBC systems have been extensively studied for their involvement in bacterial persistence, formation of persister cells, and stress adaptation [19, 20]. These systems influence growth arrest, dormancy, and antibiotic tolerance, which are crucial for bacterial survival in challenging environments [21, 22]. The VapBC family constitutes the predominant group of toxin-antitoxin loci in extreme thermoacidophiles, implying their vital significance in the physiology of these organisms. For instance, *S. tokodaii* has 25 *vapBC* toxin-antitoxin loci, while *S. solfataricus* has 22 [23]. Various *vapBC* pairs within *S. solfataricus* demonstrated differential transcriptional patterns during heat shock [17]. In vitro experiments involving a toxin responsive to heat shock, VapC6 from *S. solfataricus*, revealed its ability to target mRNA specifically. This study demonstrated VapC6 mediated targeting of transcripts coding for its cognate antitoxin *vapB6*, a transcriptional regulator (*tetR)*, and an oligo/dipeptide transport permease (*dppB-1*) [18]. Strikingly, deleting the gene encoding VapC6 from the *S. solfataricus* genome rendered the archaeon susceptible to heat shock, indicating its pivotal involvement in response to thermal stress [18]. Under uranium stress conditions, another thermoacidophile, *Metallosphaera prunae,* employs VapC toxins in post-transcriptional regulation, inducing a cellular dormant state [24]. This strategic adaptation counteracts the damage by toxic metals [24]. Moreover, previous investigations successfully identified 18 unique type II *vapBC* TA systems in *Sulfolobus acidocaldarius*, each harboring a PIN domain within its toxin protein [15]. An interesting observation from our findings was the distinct upregulation observed for at least eight specific *vapBC* pairs out of the sixteen encoded *vapBC* systems under various stressors, including heat stress, oxidative stress, and nutrient limitations [15]. This suggested their involvement in facilitating adaptive responses to individual stress types, a phenomenon termed cross-stress adaptation [15].

In the current study, we conducted a detailed biochemical and genetic analysis of the *saci_1812/saci_1813* Toxin-antitoxin pair which is upregulated under heat stress in *S. acidocaldarius* [15]. In a recent study, Lewis et al. designated this TA pair as *vapBC4* [25]. In a separate study, we simultaneously denoted the identical pair as *vapBC5*, purely based on the detection of the PIN-domain in the toxin counterpart of type II *vapBC* TA loci [15]. To maintain uniformity in the nomenclature used in the field, we uphold the identity of this TA pair as *vapBC4* in the present study. Our investigation demonstrated that the *vapBC4* TA system is a bicistronic type II toxin-antitoxin. The toxin component of this system (VapC4) exhibited a temperature-dependent ribonucleolytic activity targeted towards both mRNA and rRNA molecules *in vitro*. This enzymatic activity is believed to hinder the translation process, leading to cellular persistence efficiently. Subsequently, a deletion mutant of *vapC4* toxin resulted in decreased viability of the cells in response to heat stress at 85℃ in concordance with the observed maximal upregulation of the *vapBC4* operon in response to heat stress. This mutant also showed an increased tendency to form biofilm, suggesting a role of VapC4 toxin in biofilm formation, as was already the case for VapC14 (Saci_2183) [25].

Together, the present study demonstrated, the significance of the archaeal VapC4 toxin component within the type II TA system in promoting cellular survival under heat stress by inhibiting translation, allowing the cells to enter a persister-like state as a coping mechanism for stress.

## 2. MATERIALS AND METHODS

### Phylogenetic analysis

The SpeciesTreeAlignment.fa file generated by OrthoFinder was used as input for inference of the best amino acid substitution model and for subsequent phylogenetic inference by maximum likelihood [26]. Once we obtained the phylogenomic tree, we began searching for homologs of Saci_1812 (VapB4) and Saci_1813 (VapC4). For this, we used blastp with the parameters recommended by the developer [27]. Finally, the possible homologous proteins obtained were aligned with the MAFFT tool to identify regions at the amino acid sequence level that could be conserved [28]. The result of the phylogenomic analysis and the search for homologs was visualized through the iTOL software [29].

### Strains and growth conditions

*S. acidocaldarius* MW2000 (WT), MW1363 (Δ*vapC4*) and MW1365 (Δ*vapBC4*) and complementation strains were grown aerobically in Brock medium at pH 3, enriched with 0.1% N-Z-amine, 0.2% sucrose, and for strains not carrying a plasmid,10 µg/ml uracil, in a 75℃ incubator shaker (Thermo Scientific MaxQ 6000). Growth was tracked by assessing the optical density at 600 nm (OD_600_) using a UV-vis spectrophotometer (Dynamica HALO XB-10).

### Generation of *vapC4*, *vapBC4* knock-out strains

The plasmids pSVA6520 and pSVA6567 were generated by cloning 500 bp of the upstream and downstream regions of the target genes into plasmid pSVA431 (Table S3). The plasmids were methylated by transforming them into *E. coli* ER1821 cells harboring the additional plasmid pM.EsaBC4I (New England Biolabs, Frankfurt am Main, Germany) to evade restriction by the *Sua*I restriction system.

For generation of competent cells, *S. acidocaldarius* strains were grown in Brock medium enriched with 0.1% N-Z-amine, 0.2% dextrine, and 10 µg/ml uracil. When the optical density (OD_600_) reached 0.5–0.7, a calculated amount of the cell culture was transferred to fresh medium and harvested the following day at an OD_600_ of 0.2–0.3. The culture was cooled on ice, centrifuged for 20 min at 4000 × g at 4°C, and washed three times with 30 ml and once with 1 ml of 20 mM ice-cold sucrose. Cells were resuspended in 20 mM sucrose to a theoretical OD of 20, aliquoted into 50 μl and stored at −80°C.

Purified methylated plasmids were then used for transformation into competent MW2000 cells as described previously [30]. Colony PCR was performed to detect knock-out mutants using primers binding outside of the flanking regions. The PCR products of positive colonies were sent to sequencing analysis to confirm in frame deletion and no additional mutations.

### Complementation assays

For complementation assays, we complemented *ΔvapC4* with *vapC4* by electroporating plasmid pSVA6524 (*vapBC4* operon along with its upstream 150 bps promoter cloned into the pSVAara-FX-Stop backbone with a premature stop codon in the *vapB4* gene) into *ΔvapC4* strain. The colonies obtained after electroporation were selected on first selection plates lacking uracil. Similarly, the *ΔvapBC4+vapBC4* complementation was generated by electroporating pSVA6525 plasmid (*vapBC4* operon along with its upstream 150 bps promoter cloned into the pSVAara-FX-Stop backbone) into *ΔvapBC4* strain. Since we could not obtain the *ΔvapB4* strain, we complemented the *vapC4* gene into the *ΔvapBC4* strain, generating the *ΔvapBC4+vapC4* strain. This was performed by electroporating pSVA6524 plasmid into *ΔvapBC4* strain followed by uracil selection.

### Construction of overexpression vectors for VapB4 antitoxin and VapC4 toxin proteins

For heterologous expression of the VapB4 antitoxin and VapC4 toxin, both with a C-terminal histidine tag, the *saci_1812* (*vapB4*) and *saci_1813* (*vapC4*) genes were PCR-amplified from gDNA of *S. acidocaldarius* DSM639. The PCR amplified product was digested with dual restriction enzymes (*Nco*I and *Xho*I) and was subsequently ligated into pET28a (Kanamycin-resistant), yielding plasmid pAG153 and pAG154 for VapC4 and VapB4 respectively. The PCR product of amplified *vapB4* was additionally ligated into the multiple cloning site 2 (MCS2) of pETDuetI (Ampicillin-resistant), which harbors a C-terminal StrepII tag, constructing plasmid pAG155. For conducting expression analysis, the respective vectors were introduced into *E. coli* BL21 (DE3) cells containing the RIL Cam^r^ plasmid (Stratagene). A comprehensive list of all the strains, primers, and plasmids generated is listed in Supplementary information (**Table S1-S3**)

### Toxicity test

To conduct streak toxicity tests, cultures of *E. coli* BL21(DE3) cells carrying pAG153 and pAG154 grown overnight were streaked onto M9 minimal agar plates supplemented with kanamycin. Also, *E. coli* cells harboring both the pAG153 and pAG155 plasmids were streaked onto minimal agar plates containing a combination of both kanamycin and ampicillin. These plates were prepared with and without the addition of 0.5 mM IPTG and were incubated overnight at 37℃.

### *E. coli* growth curve analysis and cell viability assay

Overnight cultures of *E. coli* BL21(DE3) cells carrying the empty pET28a vector, pAG153, and pAG154 were diluted in 100 ml of fresh M9 minimal media. This media was supplemented with kanamycin (50 µg/ml) and 0.5mM IPTG, and the dilution was adjusted to initiate the cultures at an initial OD_600_ of 0.01. Similarly, for *E. coli* BL21(DE3) with a combination of the empty pET28a and pETDuet1 vectors, as well as a combination of pAG153 and pAG155, the overnight cultures were diluted in 100 ml of fresh M9 minimal media. This media was supplemented with kanamycin (50 µg/ml), ampicillin (100 µg/ml), and 0.5 mM IPTG. The cultures were then incubated at 37 °C. OD_600_ measurements were taken at 30 mins intervals for up to 4 hours using a UV-vis spectrophotometer (Dynamica HALO XB-10). Simultaneously, aliquots of cells were collected at each time interval for assessing cell viability using the MTT (3-(4,5-dimethylthiazol-2-yl)-2,5-diphenyltetrazolium bromide) assay. For the cell viability assessment, 10 µl of a 5 mg/ml MTT solution was added to 100 µl of cells and incubated at 37 °C for 20 mins. After incubation, the cells were centrifuged at highest speed in a benchtop centrifuge (Tarsons Spinwin MC03) for 1 min, and the supernatant was discarded. The resulting cell pellet was reconstituted in 1 ml of DMSO. Subsequently, 200 µl of the resulting purple solution was transferred to a 96-well microtiter plate, and the released purple color was measured spectroscopically at 550 nm using a Plate reader (Thermo Scientific Multiskan GO). The intensity of the purple color signifies the amount of viable cells present.

### Recombinant protein expression in *E. coli*

A 10 ml volume of an overnight grown culture of *E. coli* BL21(DE3)-RIL cells, containing the respective pAG153 and pAG154, was used to inoculate 1 L of Luria–Bertani medium supplemented with kanamycin (50 µg/ml). The cells were cultivated at 37°C until reaching an OD_600_ of 0.6. For expressing the VapC4 toxin, 150 µM IPTG (isopropyl ß-D-thiogalactopyranoside) was added, the incubation temperature was reduced to 16°C, and growth continued overnight to minimize inclusion body formation. For the VapB4 antitoxin expression, cells were induced with 500 µM IPTG and incubated at 37°C for 3-4 hours. Subsequently, the cells were collected through centrifugation (using rotor SA-300; SORVAL RC6+ Thermo-Scientific), reconstituted in lysis buffer (50 mM Tris pH 8.0, 150 mM KCl, and 10% glycerol) supplemented with the complete EDTA-free protease inhibitor cocktail (1 tablet/50 ml of lysate; Roche), frozen in liquid nitrogen, and preserved at -80 °C.

### Recombinant protein purification from *E.coli*

Before purification, frozen cell pellets were thawed on ice and added to the lysis buffer. To the lysis buffer, 1 mM lysozyme was added separately. After 30 minutes of incubation on ice, cell lysis was achieved through sonication using Soniprep 150 (DJB Labcare, UK). The resulting cell debris was eliminated by centrifugation at 15000 rpm for 30 minutes (using rotor SA-300; SORVAL RC6+ Thermo-Scientific). The supernatant was then subjected to a Ni^2+^-NTA affinity column (Qiagen) to isolate histidine-tagged proteins. The immobilized proteins underwent a stepwise wash with lysis buffer containing 10 mM and 20 mM imidazole. Depending on the specific protein, the fraction of bound proteins was then released in a lysis buffer containing 100-300 mM of imidazole. The eluted fraction was assessed by running a reducing SDS-PAGE. Subsequently, the target protein fraction was subjected to overnight dialysis in a lysis buffer (50 mM Tris pH 8.0, 150 mM KCl, and 10% glycerol). All solutions used were nuclease-free or treated with DEPC to ensure RNase free protein purification.

### Purification of single stranded 7S SRP RNA, 23S rRNA, TetR mRNA, tRNA^Met^ and 16S rRNA

The 7S SRP-RNA, 23S rRNA, TetR mRNA, Met-tRNA, and the 16S-rRNA were synthesized using in vitro transcription protocol (Thermo-Scientific). The respective genes were cloned under the T7 promoter of the pGEM3z vector, and transcription was carried out by T7 RNA polymerase using NTPs. After completion, the reaction mixture was purified by a column using RNA Purification Kit (Qiagen). A Nanodrop (Thermo Scientific, USA) quantified the purified RNA product.

### RNase Assay

Increasing concentrations of purified VapC4 toxin (Saci 1813) were incubated at 37 ℃ and 60 ℃ for 30 mins with approximately 500 ng of purified total RNA from *Sulfolobus acidocaldarius* DSM639 in a buffer containing 50 mM sodium phosphate (pH-7.0),150mM KCl and 5mM MgCl_2_. For the VapB4 neutralization assay, VapB4 antitoxin and VapC4 toxin were pre-incubated together at 75 ° C for 10 mins in a 2:1 ratio, respectively, before adding total RNA substrate, after which the reaction was held at 60 ° C for 30 mins. For determining the optimal pH for VapC4, 500 ng of total RNA from *Sulfolobus acidocaldarius DSM639* was incubated with 2µM of purified VapC4 protein in different buffers: 50 mM Tris (pH-8.0), 150 mM KCl, and 5mM MgCl_2_; 50 mM sodium phosphate (pH-6.0), 150 mM KCl, and 5 mM MgCl_2_; 50 mM citrate buffer (pH-5.0), 150 mM KCl, and 5 mM MgCl_2_; and 50 mM citrate buffer (pH-3.0), 150 mM KCl, and 5 mM MgCl_2_ at 60℃ for 30 minutes. Since the pH of a solution is dependent on temperature, the pH of the buffer at 60℃ was monitored using pH paper. In the case of in-vitro transcribed RNA substrates, 2µM of purified VapC4 was incubated with 300ng of in-vitro transcribed RNA substrates in the same buffer and incubated at 60℃ for various time intervals: 10 minutes, 20 minutes, 30 minutes, and 60 minutes. The total reaction volume was 10 µl. Each reaction was stopped by adding 2µl of 6x loading buffer and separated on 1% Tris-acetate-EDTA agarose gels with 1ug/ml ethidium bromide. All solutions used were nuclease-free or treated with DEPC.

### Co-purification assay

*E. coli* cells were co-transformed with toxin with His tag cloned into pET28a and antitoxin with StrepII tag cloned into the pETDuet1 vector. The co-transformed cells were selected onto LB agar plates containing the antibiotic kanamycin and ampicillin. Overnight primary culture at 37℃ was given with the colony obtained on the plates with both selection markers. 100 ml of secondary culture was given the following day. After the OD_600_ reached 0.6, 0.5mM IPTG induction was given for 3 hours. The cells were then harvested by centrifugation followed by lysis with a sonicator. After sonication, the cell debris was removed by centrifugation at 13000 rpm for 30 mins at 4℃. The cell lysates obtained after centrifugation were subjected to a Ni^2+^-NTA affinity column (Qiagen). The immobilized proteins underwent a stepwise wash with lysis buffer containing 10 mM and 20 mM imidazole. The fraction of bound proteins was then released in a lysis buffer containing 100-300 mM of imidazole. The eluted fraction was assessed by running a reducing SDS-PAGE followed by Western blot analysis utilizing anti-His and anti-StrepII antibodies.

### Fluorescent labeling of VapC4 and VapB4

Purified VapB4 and VapC4 were labeled with Tetramethyrhodamine (TRITC): Alexa-532 and Fluorescein isothiocyanate (FITC): Alexa-488 fluorescent probes, respectively, following the manufacturer’s instructions (Sigma-Aldrich, USA). In brief, VapC4 (50 µM) and VapB4 (75 µM) in 50 mM Na-Phosphate Buffer supplemented with 100 mM Sodium bicarbonate (pH 9.0) were combined with FITC or TRITC fluorescent probes, respectively. FITC and TRITC stock solutions were prepared in anhydrous DMSO at 600 µg/ml concentrations. For labeling the protein samples, 50 µl of the FITC or TRITC fluorescent probe was taken from the stock solution and gradually introduced into 1 ml of the protein. The protein sample was gently and continuously stirred for 2 hours at room temperature. A Bio-Gel A column (Bio-Rad) was utilized to segregate the labeled protein from the unbound probe. The collected fraction containing the labeled VapB4 or VapC4 was dialyzed against a phosphate buffer (50 mM, pH 7.2) at room temperature for 24 hours. The evaluation of the protein labeled with FITC or TRITC involved measuring the absorbance of the protein at 280 nm and the corresponding fluorescence probes at 495 nm (for FITC) or 542 nm (for TRITC). The extent of FITC or TRITC labeling on the protein was determined based on the F/P ratio, which was approximately 0.5 for both probes, confirming optimal protein labeling. Lastly, the labeled proteins were verified by running an SDS-PAGE and scanning using a Typhoon scanner (GE Healthcare, Danderyd, Sweden).

### Fluorescence resonance energy transfer (FRET)

The fluorescence resonance energy transfer (FRET) method was employed to ascertain the interaction between VapC4 toxin and VapB4 antitoxin proteins. FITC-labeled VapC4 (100 nM), serving as the donor, was titrated with TRITC-labeled VapB4 (25-2000 nM) acting as acceptor. A no FRET control set was also set where FITC-labeled VapC4 (100 nM), serving as the donor, was titrated with only buffer. The reaction mixture was excited at 495 nm, and the fluorescence emission spectra were taken from 505-625 nm at 60°C using a JASCO f-8500 spectrophotometer (JASCO International Co., Ltd., Japan). Subsequently, the donor and acceptor emission spectra were deconvoluted using Igor pro 6 (WaveMetrics, Portland, Oregon, USA). The emission intensity of FITC at 520 nm was individually measured during various titrations and used to compute the dissociation rate constant (Kd) using the subsequent equation:

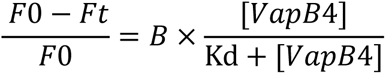

Where Ft represents fluorescence intensity at 520 nm for various concentrations of VapB4, and F0 is the fluorescent intensity at [VapB4] = 0. Kd denotes the dissociation rate constant, and B represents the maximum intensity reached at the saturation of binding.

### *S. acidocaldarius* growth curve analysis

Exponentially growing cultures of *S. acidocaldarius* MW2000 (WT), Δ*vapC4*, and Δ*vapBC4* were diluted into 50 ml of Brock medium at pH 3, supplemented with 0.1% N-Z-amine, 0.2% sucrose, and 10 µg/ml uracil. The dilution was adjusted to an initial OD_600_ of 0.01. The cultures were then incubated in a 75℃ incubator shaker (Thermo Scientific MaxQ 6000). The growth progression of the organisms was checked at four-hour intervals by measuring the optical density at 600 nm (OD_600_) using a UV-vis spectrophotometer (Dynamica HALO XB-10). The experiment was performed in three biological replicates.

To assess growth under heat stress, cell cultures in the exponential growth phase were diluted to an OD_600_ of 0.5. Subsequently, 1 ml of the diluted cells was dispensed into 2 ml microcentrifuge tubes and subjected to heat stress at 85°C in a shaking dry heat block. Growth progression was tracked by recording the OD_600_ at hourly intervals. As a control, the diluted cell cultures were maintained at 75°C in a shaking dry heat block. The experiment was performed in three biological replicates.

### Cell viability assay for *S. acidocaldarius*

To evaluate the ability of the generated *Sulfolobus* strains to survive under heat stress, an MTT assay was conducted at various time points during heat stress exposure. All knock-out and complementation strains were cultured until the mid-exponential phase (OD_600_ 0.5). Subsequently, 2 ml of culture from each strain was transferred to microcentrifuge tubes. These tubes were then exposed to heat stress by incubating at 85°C in a Thermo Shaker incubator (BenchTop Lab Systems BT-MTH-100) for different durations (15, 30, 45, and 60 mins), with strains maintained at 75℃ serving as controls. The heat stress temperature was determined following a previous study by Baes et al. [31]. After incubation, 100 µl of the cultures were mixed with 10 µl of MTT reagent (5 mg/ml) and incubated for 1 hour at 75°C in an incubator (Thermo Scientific MaxQ 6000). Following centrifugation at 13,000 rpm using a benchtop centrifuge (Tarsons Spinwin MC03) for 1 min, the supernatant was discarded, and the pellets were dissolved in 1 ml DMSO. Subsequently, 200 µl of the resulting purple solution was transferred to a 96-well microtiter plate, and the released purple color was measured spectroscopically at 550 nm using a Plate reader (Thermo Scientific Multiskan GO). The intensity of the purple color signifies the amount of viable cells present. To calculate the percentage survivability based on the MTT assay absorbance at 550 nm, the following formula was used:

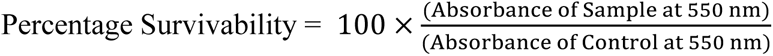

Where the “Absorbance of Sample” refers to the absorbance measured for each time point after heat stress, and the “Absorbance of Control” is the absorbance of the corresponding control set (not subjected to heat stress). The experiment was performed in three biological replicates.

### Biofilm formation

All knock-out and complementation strains were grown to mid-exponential phase (OD_600_ 0.5). Subsequently, cells were inoculated into 24-well microtiter plates filled with Brock medium at pH 3, enriched with 0.1% N-Z-amine, 0.2% sucrose, and 10 µg/ml uracil (for WT, Δ*vapC4*, and Δ*vapBC4*), or only 0.1% N-Z-amine and 0.2% sucrose (for the complementation strains). The cells were inoculated into 24 well plates in such a way such that their initial OD_600_ was 0.01. For the heat stress-induced biofilm formation, the cells were initially subjected to heat stress at 85℃ for different durations (15 mins, 30 mins, 45 mins, and 60 mins). After each heat stress period, the cells were collected and subjected to an MTT assay to quantify viable cell counts. Following the assay, an equal number of heat-stressed viable cells, determined from the MTT results, were inoculated into each well of the 24-well plates. The 24-well plate was sealed tightly with a gas-permeable sealing membrane (Breathe-Easy, Diversified Biotech, Boston, MA, USA) and incubated at 75 °C in an incubator (Thermo Scientific MaxQ 6000) for 48 hours within a humidity chamber to avoid evaporation. The initial OD_600_ was adjusted to 0.01, following the procedure outlined by Koerdt et al. [32]. After 48-hour incubation, the 24-well microtiter plates were cooled, and 200 µl of vegetative cells from each sample were transferred to a new 96-well microtiter plate. The OD_600_ of the supernatant containing the vegetative cells was determined using a 96-well plate reader (Thermo Scientific MultiSkan GO). The remaining supernatants were carefully discarded without disturbing the biofilm adhered to the 96-well microtiter plate. Subsequently, 200 µl of a 0.5% w/v crystal violet (CV) solution was added to each well and incubated for 10 minutes at room temperature. Once again, the CV supernatant was removed, and the biofilm adhered to the well plate was washed three times with Brock medium at pH 5 until residual crystal violet was eliminated. One ml of 30% acetic acid was added to each well to release the crystal violet absorbed by the biofilm-forming cells, and absorbance was measured at 570 nm. The OD_570_/OD_600_ correlation index was employed to assess the efficiency of biofilm formation. The experiment was conducted in biological replicates.

### Statistical Analysis

GraphPad Prism 6 was utilized for statistical analyses. The values in the figures represent the mean of three replicates ± the standard error of the mean. We used two-way ANOVA and the Bonferroni post-test for the statistical data comparison. In all analyses, a significance level of p < 0.05 was considered.

## 3. RESULTS

### The *vapB4* and *vapC4* genes are co-transcribed in *S. acidocaldarius*

The *Sulfolobus acidocaldarius* DSM 639 genome contains a potential *vapBC4* operon (*saci_1812* and *saci_1813*, depicted in **Figure 1a**). Subsequent evaluation of the toxin gene sequence encoded a VapC-like member of the PIN domain superfamily of ribonucleases. The identified antitoxin gene encoded a CopG family transcriptional regulator, possessing an RHH-like domain usually observed in VapB antitoxins. Notably, sequence analyses showed an overlap for the translational start codon of *vapC4* with the translational stop codon of *vapB4*, strongly implying translational coupling (**Figure 1a**). To confirm the existence of these two genes as a bicistronic operon, reverse transcriptase (RT)-PCR experiments were conducted. The primers selected targeted a segment that spanned from the end of the antitoxin gene to the start of the toxin gene. As shown (**Figure 1b**), these genes are indeed expressed as a single transcriptional unit. Since most of the TA systems are inherited through horizontal gene transfer, we sought to assess the conservation of the VapBC4 TA system beyond *Sulfolobus* species. We searched for homologous genes in a set of 78 archaeal genomes (Supplementary Information **Table S4**). We identified 25 pairs of toxin-antitoxin genes in various lineages, including the superphylum Asgardarchaeota, DPANN, TACK, and the phylum Euryarchaeota. The specific organisms harboring these potential homologs are detailed in Supplementary Information (**Table S5** and **Figures S1-S2)**. None of the listed VapBCs have been functionally characterized to date. This distribution suggests that the VapBC4 TA system, or its homologs, are not confined to a specific taxonomic group but are instead widely distributed among archaeal taxa, suggesting their significance in archaeal biology, and highlighting the need for further investigation into their roles and evolutionary dynamics.

**Figure 1:**
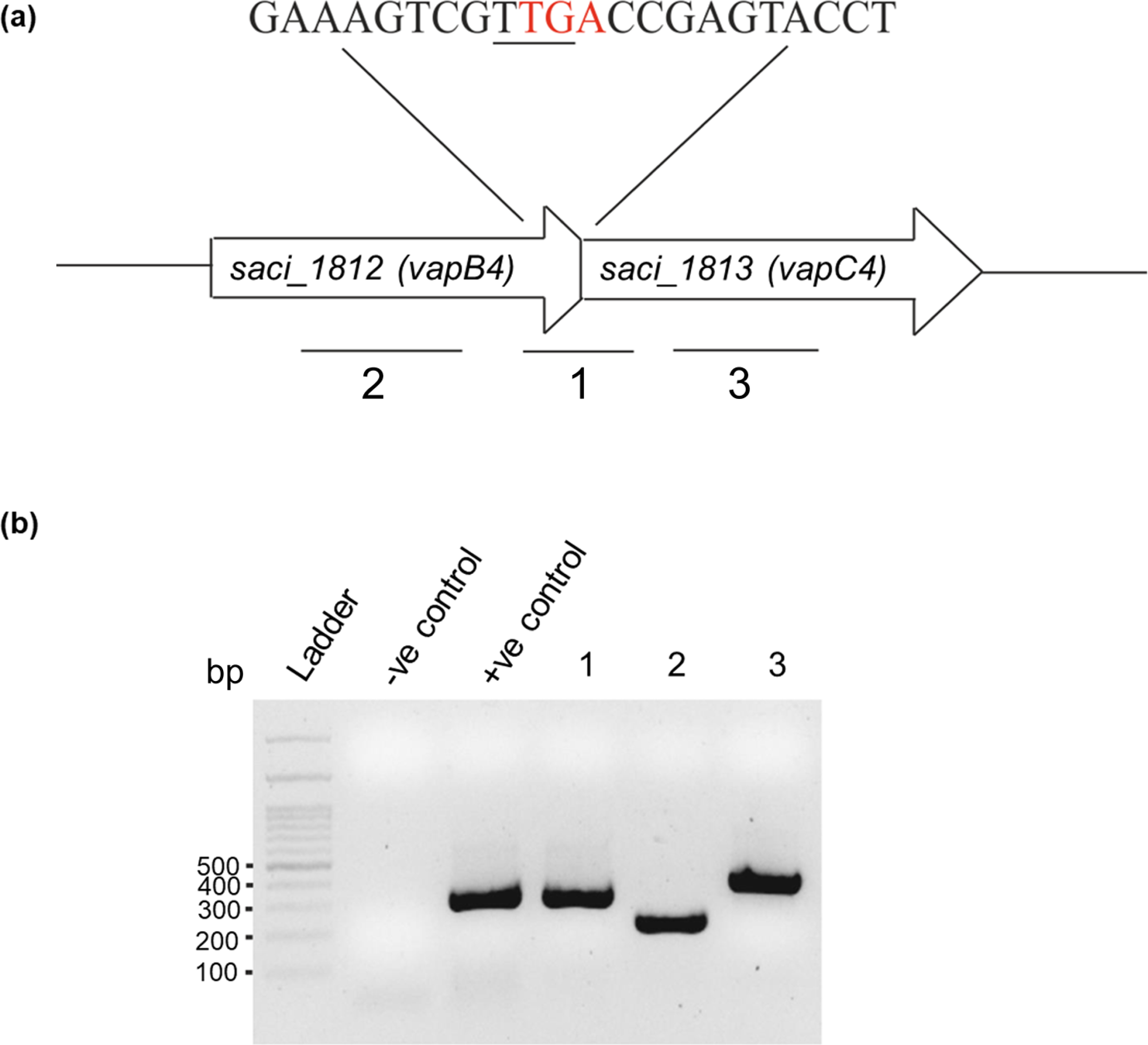
Genomic organization of *vapBC4* Toxin-antitoxin gene locus in *Sulfolobus acidocaldarius*. a) Schematic representation of the genes *saci_1812* (VapB4 antitoxin) and *saci_1813* (VapC4 toxin). The stop codon (TGA) of the *vapB4* gene (marked in red) overlaps the start codon (TTG) of the *vapC4* gene (underlined). b) Agarose gel showing PCR amplification of the respective region of the operon marked as 1, 2 and 3 in a), using cDNA as template (1-intermediate region of the *saci_1812* (*vapB4*) antitoxin and *saci_1813* (*vapC4*) toxin gene; 2-region of the *saci_1812* (*vapB4*) antitoxin; 3-region of the *saci_1813* (*vapC4*) toxin. Negative control with no reverse transcriptase was used to verify that RNA samples do not contain gDNA contamination. In the positive control, gDNA was used as a template to amplify the intermediate portion of the *saci_1812* and *saci_1813* gene marked as “1”.

### Heterologous expression of VapC4 leads to cell growth inhibition in *E. coli*

*E. coli* cells harboring different constructs, including the empty pET28a vector, pAG153 (VapC4 toxin) and pAG154 (vapB4 antitoxin), and the combination of both pAG153 and pAG155 (vapB4 antitoxin), were streaked onto M9 minimal media agar plates supplemented with the appropriate antibiotics for selection and 0.5 mM of IPTG to induce gene expression. Cells expressing the VapC4 toxin exhibited growth inhibition in comparison to cells expressing the VapB4 antitoxin or carrying the empty vector (**Figure 2a, upper row**). Conversely, the co-expression of the VapB4 antitoxin and the VapC4 toxin rescued the cell growth (**Figure 2a, lower row**). A growth curve analysis of bacterial cells expressing the different genes was performed to further investigate this impaired growth phenomenon upon toxin expression and the alleviation of toxin activity upon antitoxin expression. The VapC4 toxin-expressing cells showed significant growth defects compared to those carrying empty pET28a and VapB4 antitoxin expressing plasmid (**Figure 2b**). The growth was partially rescued when the VapB4 antitoxin was co-expressed along with the VapC4 toxin (**Figure 2b**). A similar observation was noted in bacteria when VapC toxins from various sources, including a plant pathogen *Acidovorax citrulli*, a deep-sea marine bacterium *Streptomyces sp.* SCSIO02999, and a human pathogen non-typeable *Haemophilus influenzae*, were expressed heterologously in *E. coli* cells [33–35]. It is particularly surprising to observe that the toxin derived from a thermoacidophilic archaeon can effectively induce bacteriostasis in the mesophilic *E. coli* cells.

**Figure 2:**
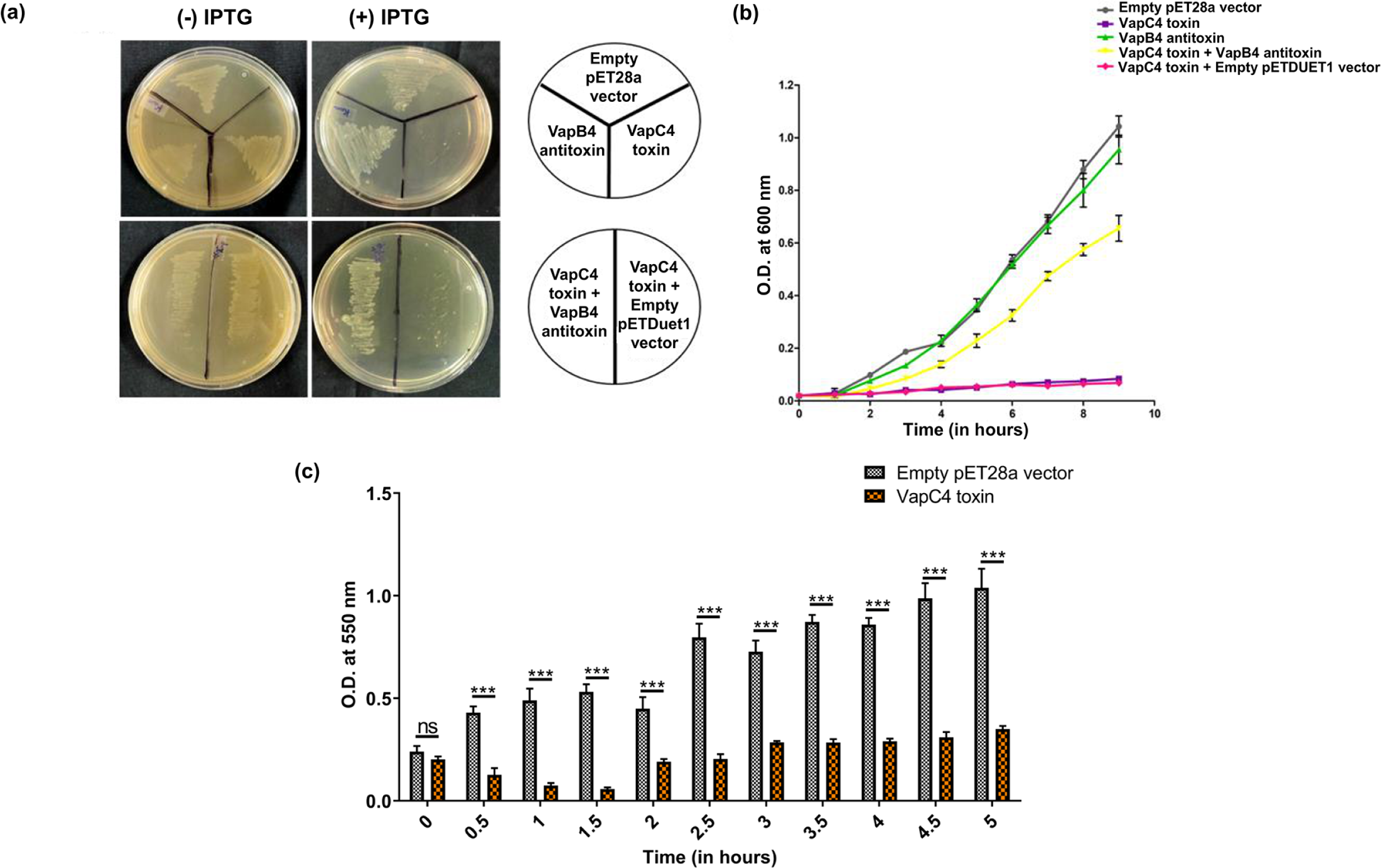
The heterologous expression of VapC4 toxin (Saci_1813) inhibits growth in-vivo leading to persister cells. (a) *E. coli* streaked M9 minimal media plates expressing VapC4 toxin, VapB4 antitoxin, empty pET28a vector, VapC4 toxin + VapB4 antitoxin and VapC4 toxin + empty pETDuet1 vector (b) Growth profiles of *E. coli* CBL21(DE3) cells expressing the empty pET28a vector, VapC4 toxin, VapB4 antitoxin, both VapC4 toxin + VapB4 antitoxin as well as VapC4 toxin + empty pETDuet1 vector in M9 minimal media containing 0.5mM IPTG. c) Bar graph showing the viability of cells expressing empty pET28a vector and VapC4 toxin with increasing time interval. Experiments were performed in triplicates. The statistical significance, P < 0.05, is indicated as *, P < 0.01 is indicated as **, and P < 0.001 is indicated as ***; n.s.denotes not significant.

### VapC4 toxin expression in *E. coli* leads to persister cells

Previous reports in bacteria have indicated that the expression of toxins can induce alterations in cellular metabolism, potentially triggering the emergence of the persister phase or leading to cell death [36]. The cells’ fate depends on the specific nature of stress and the type of toxin in play. To understand what is happening to the *E. coli* cells that result in the growth impairment caused by the *Saci* VapC4 toxin expression, we conducted a MTT assay to assess cell viability. We aimed to investigate the fate of *E. coli* cells upon the induction of the VapC4 toxin. In the cell viability test, the absorbance at 550 nm displayed a gradual increase for cells containing the empty pET28a vector (control), showing no influence on their growth. In contrast, following 1 hour of induction, a reduction in absorbance (OD_550_) became apparent, signifying that upon VapC4 overexpression, a portion of the cells dies (**Figure 2c**). After 2 hours, cell growth was gradually restored and remained stable. This pattern indicates that the overexpression of VapC4 generates a bacteriostatic effect where the cells are viable but incapable of dividing. This phenomenon was also observed when *Leptospira* VapC and *Mycobacteria* VapC46 were expressed in *E. coli* [10, 37].

### The RNase activity of VapC4 from *S. acidocaldarius* is catalyzed by high temperatures and is effectively counteracted by the inhibitory action of VapB4

The purification of the VapC4 toxin protein from *E. coli* presented a significant challenge due to the observed inhibition of cell growth upon protein overexpression. In an attempt to overcome this hurdle, VapC4 expression was gradually induced at a very low temperature of 16℃ and different IPTG concentrations were tested. Surprisingly, at an IPTG concentration of 0.1 mM, the cells could express the toxic protein without causing any detrimental impact on their growth (as depicted in **Figure 3a**). This successful outcome was corroborated by SDS-PAGE analysis, which indicated the presence of VapC4 protein with a migration pattern aligning with that of the 16 kDa marker (as illustrated in **Figure 3b**). Conversely, the VapB antitoxin was produced in *E. coli* growing at 37℃ at and induced with 0.5 mM IPTG for 3-4 hours. Subsequent SDS-PAGE analysis confirmed the presence of VapB4 protein whose migration aligned with that of the 9 kDa marker (as represented in **Figure 3c**).

**Figure 3:**
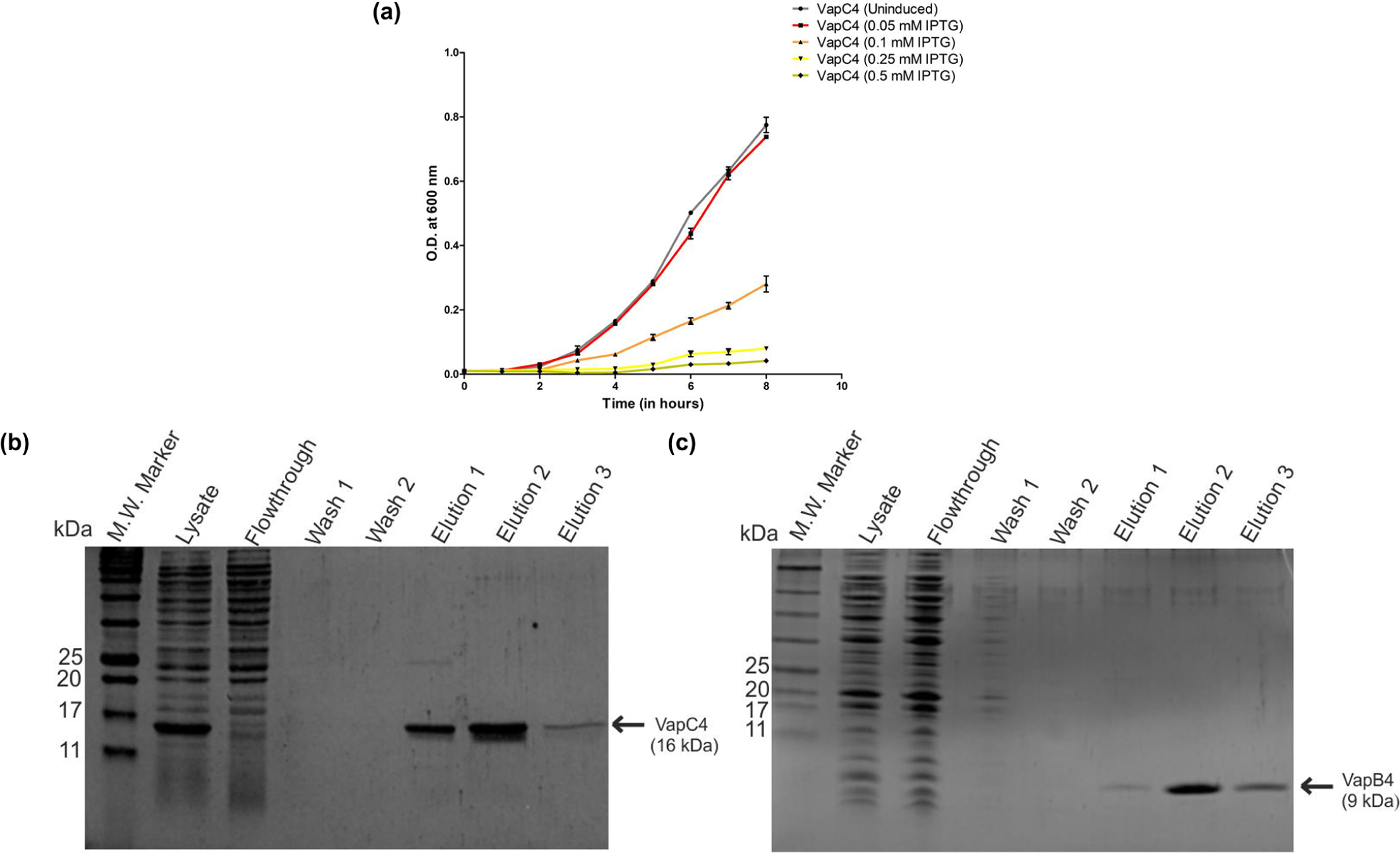
Purification of VapC4 toxin and vapB4 antitoxin. a) Growth profile of *E.coli* BL21(DE3)-RIL cells expressing VapC4 toxin in the presence of different IPTG concentrations at 16 ℃ induction temperature. b) A 15% SDS PAGE shows the successful purification of VapC4 toxin using Ni^2+^-NTA affinity chromatography around 16 kDa. c) A 15% SDS PAGE shows successful purification of VapB4 antitoxin using Ni^2+^-NTA affinity chromatography around 9 kDa.

The VapC proteins are known for their ribonucleolytic activity due to the presence of the characteristic PIN domain in their structure [38]. When the purified VapC4 toxin was incubated with total RNA extracted from *S. acidocaldarius* at two different temperatures, 37℃ and 60℃, it was observed that VapC4 was able to degrade the RNA at both temperatures. Complete degradation of RNA was observed with 1.3 µM of VapC4 at 60℃ compared to that at 37℃, aligning with the fact that *S. acidocaldarius* is a thermophilic organism. (**Figure 4a-b**). A similar observation was made with the *S. solfataricus* VapC6 toxin, demonstrating more efficient cleavage of total RNA at 80°C in contrast to temperatures of 37°C and 60°C [18]. As the *vapC4* toxin gene was identified within a bicistronic operon alongside the *vapB4* antitoxin gene, a characteristic feature of type II Toxin-antitoxin systems, we proceeded to investigate the inhibitory function of the cognate VapB4 antitoxin on the VapC4 toxin. In this case, purified VapB4 antitoxin was incubated with the VapC4 toxin at two different temperatures, 37℃ and 60℃. At 60℃ the RNase activity of VapC4 was substantially inhibited compared to the activity observed at 37℃ (as illustrated in **Figure 4c**). This observation validates the protective role of the VapB4 antitoxin against VapC’s action on substrates. Additionally, it confirms that only the VapC4 toxin is responsible for RNase activity, ruling out the involvement of any other potential RNase from *E. coli*.

**Figure 4:**
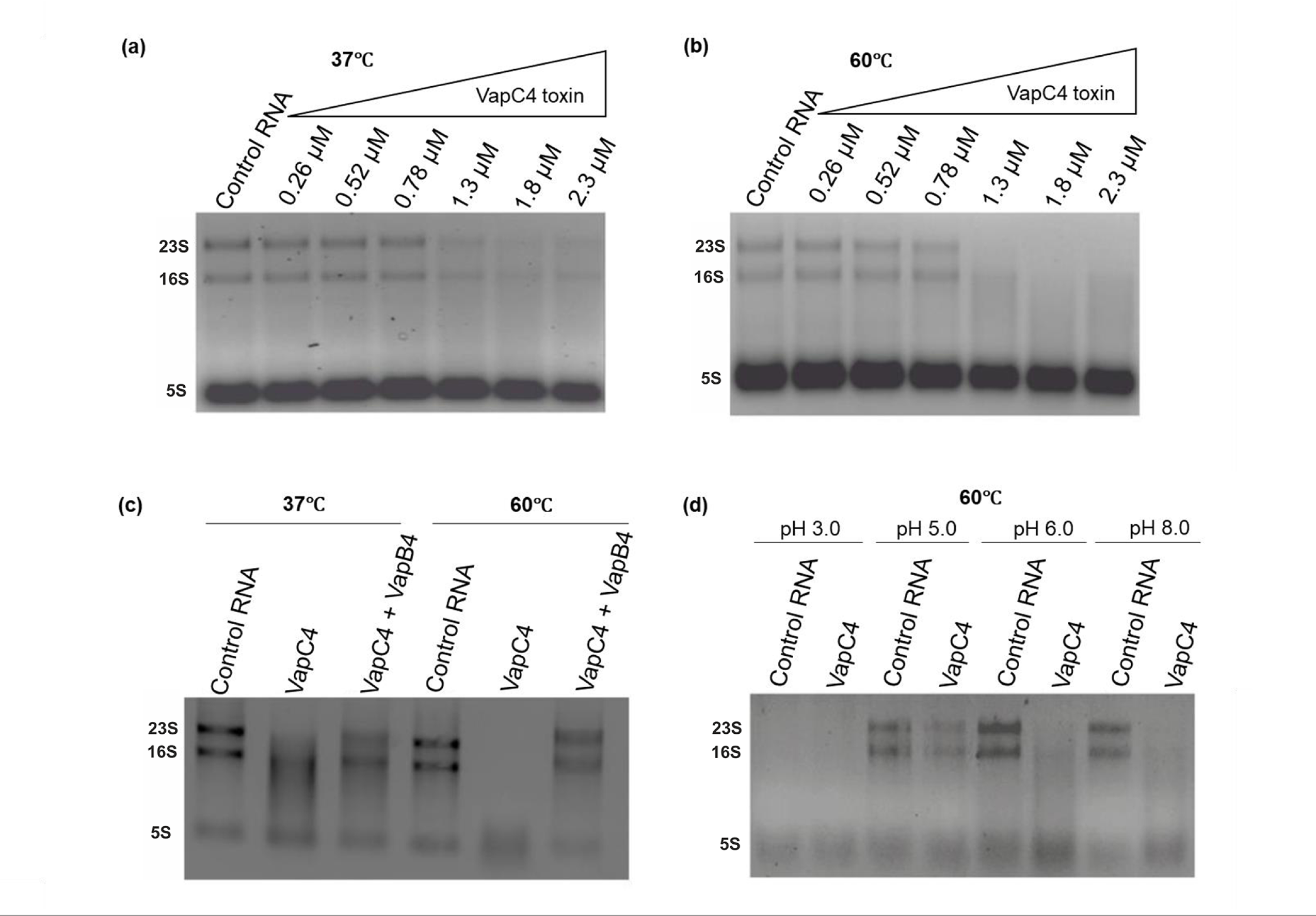
Ribonucleolytic Activity of VapC4 Toxin. 1% agarose gel demonstrating RNase activity of VapC4 toxin at two distinct temperatures: a) 37℃ and b) 60℃ on *S. acidocaldarius* total RNA. c) 1% agarose gel illustrating the inhibitory effect of VapB4 antitoxin on VapC4 toxin at two different temperatures, 37℃ and 60℃. d) Agarose gel displaying RNase activity of VapC4 at different pH at 60℃.

An RNase assay was conducted to determine the optimal pH conditions under which the VapC4 toxin exhibits its highest efficacy. The VapC4 protein was subjected to incubation with total RNA from *S. acidocaldarius* at different pH values, all maintained at a temperature of 60℃. As expected, VapC4 demonstrated enhanced RNA hydrolysis efficiency across the pH range of 6 to 8 (**Figure 4d**). This observation aligns with expectations, considering the internal pH of *S. acidocaldarius* is around 6.5 [39].

### VapC4 toxin can hydrolyze mRNA and rRNA in vitro

A prior study proposed that VapC1 and VapC2 derived from non-typeable *Haemophilus influenzae* (NTHi) can inhibit translation by cleaving initiator tRNA^fMet^, thereby inducing the formation of dormant cells [40]. However, VapC6 from *S. solfataricus* has been shown to cleave *dppB-1*, *tetR*, and *vapB6* mRNA [18]. Given that VapC exhibits varying RNA substrate specificity depending on the organism, and with the confirmed RNAse activity of the VapC4 toxin, we wanted to identify its specific substrate. We synthesized different classes of RNA substrates from *S. acidocaldarius* and evaluated the VapC4 RNase activity on these transcribed RNA molecules to determine the substrate specificity (**Figure 5a**). To this end, the purified VapC4 toxin was incubated with various in-vitro transcribed RNAs –7S SRP RNA, tRNA^Met^, 16S rRNA, 23S rRNA, and TetR mRNA-at 60℃. Notably, the VapC4 cleaved the rRNA and mRNA, while it did not hydrolyse the 7S SRP RNA and tRNA^Met^ (**Figure 5b-f**). Interestingly, while SRP RNA and tRNA^Met^ remained unaffected, the complete degradation of 23S rRNA and TetR mRNA required 10 mins, whereas 16S rRNA took 20 mins for complete breakdown (**Figure 5b-f**). Hence, the VapC4 enzyme from *S. acidocaldarius* can degrade mRNA and rRNA *in vitro*. It can be speculated that the significance of mRNA hydrolysis lies in the direct inhibition of translation, while its rRNA hydrolysis indirectly inhibits translation. As published previously, the cleavage of 16S rRNA causes a separation of the anti-Shine-Dalgarno sequence from the ribosome [41]. Consequently, the ribosome’s ability to scan the initiation codon of the mRNA is impaired, leading to a subsequent hindrance in protein translation. On the other hand, the cleavage of the 23S rRNA triggers the disassembly of the 50S and 30S ribosomal subunits, thereby compromising the entire translation apparatus [42]. This suggested that VapC4 might serve as a dual substrate-specific RNase, targeting both mRNA and rRNA for degradation, ultimately leading to the inhibition of translation. Previous studies on *Metallosphaera* VapCs demonstrated that VapC4 specifically recognized a GAAG consensus motif, whereas VapC7 and VapC8 targeted the degenerate consensus motifs (A/U)AG(G)A and (U/G)AAU, respectively, within the 16S and 23S rRNA [24].

**Figure 5:**
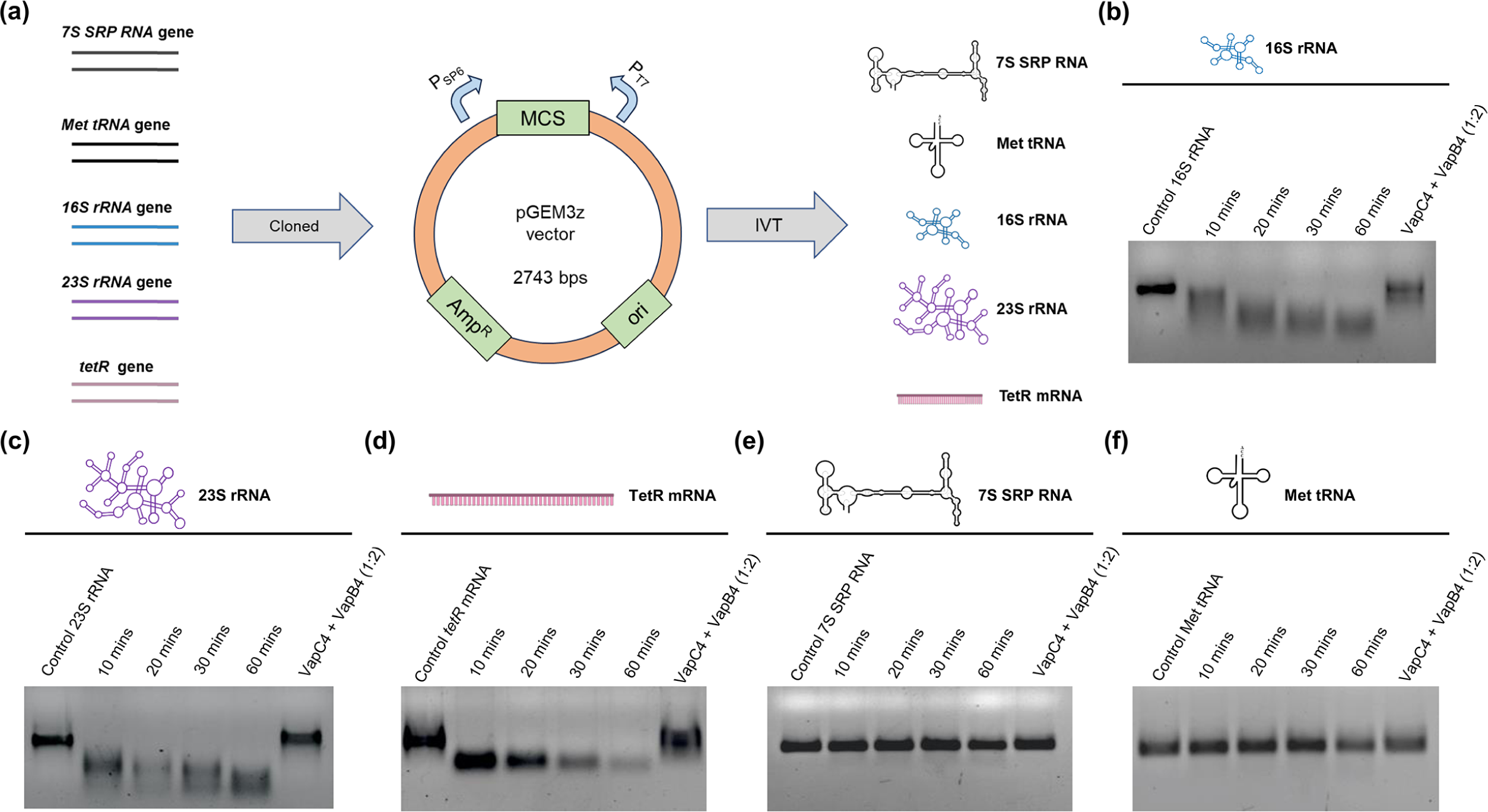
RNA specificity of VapC4 Toxin. a) Schematic showing different classes of single-stranded RNA substrate generation by In-vitro-transcription (IVT) using pGEM3z vector (P_SP6_ and P_T7_ denotes SP6 and T7 promoter respectively). The RNase activity of the VapC4 toxin was examined at 60℃ using various classes of RNA substrates from *S. acidocaldarius*, prepared through In-vitro transcription (IVT) reactions: b) 16S rRNA c) 23S rRNA d) TetR mRNA e) 7S SRP RNA, and f) tRNA^Met^.

### VapB4 antitoxin interacts with VapC4 toxin to inhibit its effect

To ascertain whether the inhibition caused by the VapB4 antitoxin occurs through the binding of the antitoxin to the VapC4 toxin rather than the toxin’s target (RNA), we performed toxin pull-down experiment. To that end, both the VapC4 toxin with a His-tag and the VapB4 antitoxin with a StrepII tag were expressed in *E. coli*. These lysates were subjected to Ni-NTA affinity chromatography, which resulted in the elution of both proteins combined within a single (elution 2) fraction (**Figure 6a**). The size of the eluted proteins corresponded to the migration pattern of both the VapC4 toxin and the VapB4 antitoxin. This co-purification strongly indicates the formation of a complex between the toxin and antitoxin within the cellular environment. This complex formation further suggests that the antitoxin hinders the toxin’s activity by strongly binding to the toxin’s active site, fulfilling the type II TA system criterion. A control set was also prepared using cell lysate from *E. coli* cells that expressed only VapB4 antitoxin with a StrepII tag. When this lysate was subjected to Ni-NTA affinity chromatography, no band corresponding to the VapB4 antitoxin was observed in the elution 2 fraction (**Figure S3**). Verification through Western Blot analysis, using anti-His antibodies and anti-Strep-II antibodies, confirmed that the observed protein bands indeed corresponded to the VapC4 toxin and VapB4 antitoxin, respectively (**Figure 6b**).

**Figure 6:**
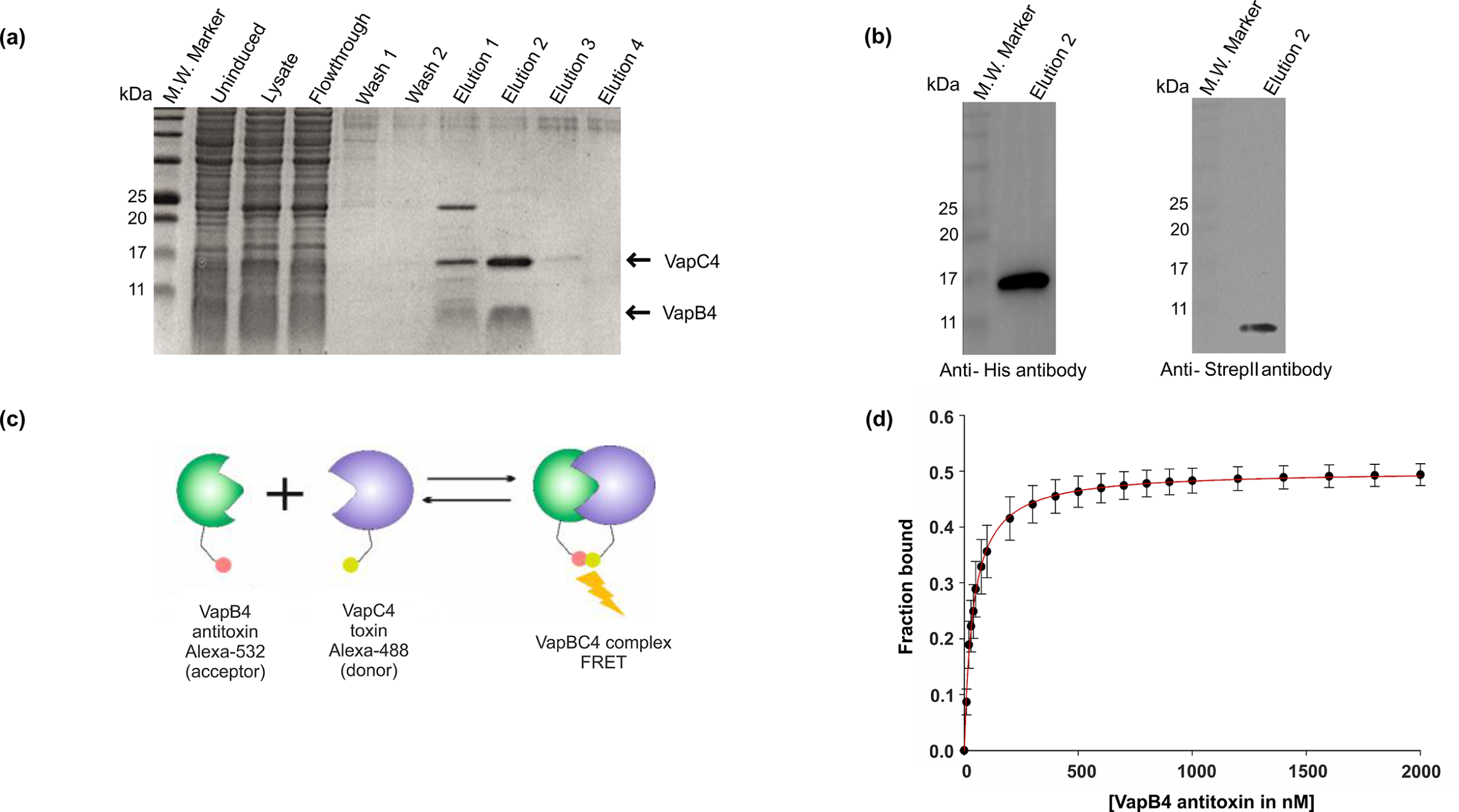
Interaction between VapC4 toxin and VapB4 antitoxin. a) 15% SDS-PAGE showing co-elution of VapC4 toxin (at 16 kDa position) along with VapB4 antitoxin (at 9 kDa position) in the Elution 2 fraction. b) Western Blot analysis with anti-His and anti-Strep-II antibodies confirmed the presence of VapC4 toxin and VapB4 antitoxin, respectively. c) VapC4 toxin tagged with Alexa-488 (FITC) was titrated with increasing concentration of VapB4 antitoxin tagged with Alexa-532 (TRITC), resulting in FRET between TRITC (acceptor) and FITC (donor). d) Quantification of the binding of VapB4 antitoxin to VapC4 toxin using the equation outlined in the Material and Methods section revealed a dissociation constant (Kd) of 40 ± 2.2 nM, indicating a strong and reliable binding affinity.

To gain a quantitative understanding of the interaction between the VapC4 toxin and VapB4 antitoxin, a Fluorescent Resonance Energy Transfer (FRET) analysis was conducted. In this experiment, the VapB4 antitoxin was labeled with the fluorescent dye Alexa-532 (TRITC), while the VapC4 toxin was labeled with the fluorescent dye Alexa-488 (FITC) (**Figure 6c**). As the concentration of the VapB4 antitoxin was raised, an increase in energy transfer was detected. The dataset was subjected to fitting using the binding equation outlined in the Materials and Methods section (**Figure 6d**). The resulting dissociation constant (K_d_) derived from the experiment was determined to be 40 ± 2 nM, highlighting a robust interaction between the VapC4 toxin and VapB4 antitoxin.

### VapC4 toxin enables the cells to withstand heat stress

The maximal upregulation of the *vapBC4* operon during heat stress suggested an important role in heat stress adaptation [15]. In the present study, we generated the *S. acidocaldarius* deletion strains Δ*vapC4,* and Δ*vapBC4* and subjected them to heat stress at 85℃ for different time intervals and then determined the viability of the cells for each time point using an MTT assay. Deletion strains Δ*vapC4* and Δ*vapBC4* showed no difference in growth at 75℃ compared to the WT (**Figure 7a**). However, the *vapC4* toxin complementation in the *vapBC4* background strain led to a slower growth probably due to the toxic nature of VapC4 (**Figure 7a**).

**Figure 7.**
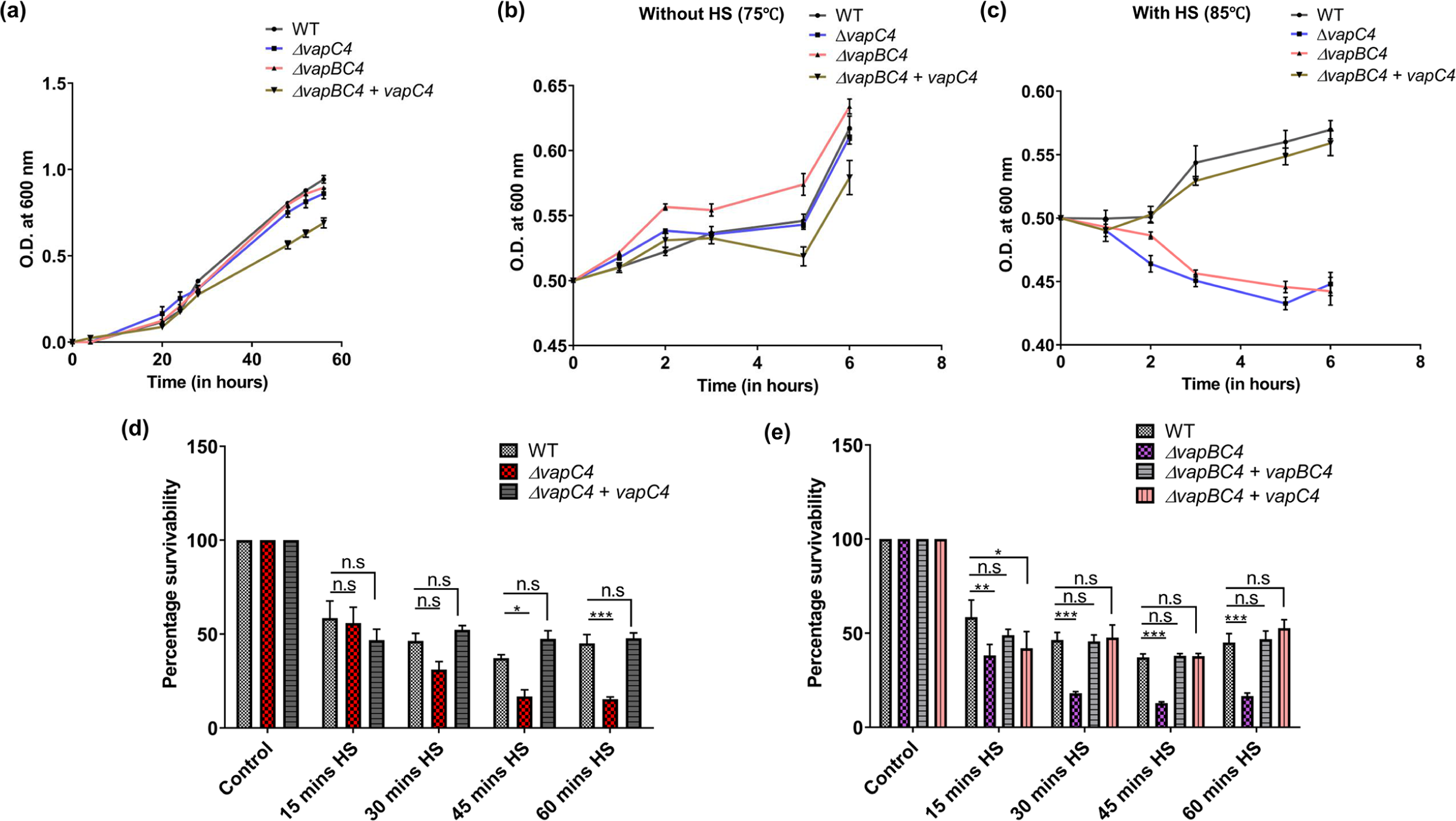
Impact of VapC4 Toxin on Heat Stress Survival. a) Growth curve analysis of all the Knock-out strains and *vapBC4* + *vapC4* at 75 ℃ in Brock medium at pH 3, supplemented with 0.1% N-Z-amine,0.2% sucrose and 10 µg/ml uracil (if required). b) Growth curve analysis of all the Knock-out strains and *vapBC4* + *vapC4* complementation strains at 75 ℃ (optimum growth condition) from a starting OD_600_ 0.5 measured for 6 hours. c) Growth curve analysis of all the Knock-out strains and *vapBC4* + *vapC4* complementation cell at 85℃ (HS-heat stress) from a starting OD_600_ 0.5 measured for total duration of 6 hours. d) Graph showing percentage survivability of *ΔvapC4* cells vs. WT during HS at 85 ℃ at different time intervals: Control (5 mins before HS), 15 mins HS, 30 mins HS, 45 mins HS, and 60 mins HS. The *ΔvapC4* + *vapC4* indicates the *vapC4* complementation strain e) Graph showing percentage survivability of *ΔvapBC4* cells vs. WT and *ΔvapBC4* + *vapC4* vs. WT during HS at 85 ℃ at different time intervals: Control (5 mins before HS), 15 mins HS, 30 mins HS, 45 mins HS and 60 mins HS. The *ΔvapBC4* + *vapBC4* indicates the *vapBC4* complementation strain.

Furthermore, upon subjecting cells in the mid-log phase (OD_600_ 0.5) to heat stress at 85℃, a noticeable growth impairment was evident in the *ΔvapC4* and *ΔvapBC4* strains, characterized by an inability to divide compared to cells cultivated at the optimal growth temperature of 75℃ (**Figure 7b-c**). Notably, despite exhibiting less efficient growth under heat stress conditions compared to their optimal growth environment, both the wild-type (WT) and *ΔvapBC4* + *vapC4* complemented cells showed resilience to heat shock (**Figure 7b-c**). As OD values can include readings from both living and dead cells, we sought to specifically assess the quantity of viable cells following heat stress. To achieve this, we conducted an MTT assay for a period of one hour of heat stress. Using the MTT assay, we found that the Δ*vapC4* cells could not withstand heat stress (HS) compared to WT cells, and the Δ*vapC4* complemented with *vapC4* (**Figure 7d**). The same decrease in the percentage survivability was also observed for Δ*vapBC4* cells (**Figure 7e**). Since we were unable to generate a *ΔvapB4* strain, we tried to complement the *vapC4* gene into Δ*vapBC4* background (Δ*vapBC4*+*vapC4*) to check whether the toxic effect of VapC4 is indeed responsible for rescuing the cells during heat stress. As expected, Δ*vapBC4*+*vapC4* cells were resilient to heat stress at 85℃ (**Figure 7e**). Therefore, our findings suggest that VapC4 might help *S. acidocaldarius* form persister-like cells that can withstand heat stress.

### The *vapC4* toxin gene deletion aggravates biofilm formation

Biofilm formation is a common survival strategy adopted by different bacteria and archaea in response to environmental stresses [43–45]. Biofilms offer protection against adverse environmental conditions, enhance nutrient availability, and facilitate genetic exchange [46]. Understanding the relationship between VapBC4 and biofilm formation would show how this TA system contributes to stress resistance and survival strategies in these thermoacidophiles. We conducted a biofilm formation assay using the crystal violet staining method with the *S. acidocaldarius* strains cultivated in 24-well microtiter plates for 48 hours. It was observed that the Δ*vapC4* cells exhibited increased biofilm formation compared to WT and *ΔvapC4+vapC4* complementation strains (**Figure 8a**). A significant increase in biofilm formation was also observed for Δ*vapBC4* cells compared to WT and *ΔvapBC4+vapBC4* complementation (**Figure 8b**). Further complementation of the *vapC4* gene into the Δ*vapBC4* background led to decreased biofilm formation (**Figure 8b**). Hence, this study revealed that the VapC4 toxin plays an inhibitory role in forming biofilms. In a previous study, a comparable phenomenon of biofilm overproduction was noted in both the *ΔvapC14* and *ΔvapBC14* strains, where it was hypothesized that the VapC14 (Saci_2183) toxin targets crucial biofilm-related RNAs, including the transcript of the recognized Lrs14-like biofilm activator (*saci_1223*) in *S. acidocaldarius* [25, 47]. We also investigated the impact of heat stress for various time spans (15 mins, 30 mins, 45 mins, and 60 mins) on the biofilm formation of *S. acidocaldarius*. It was observed that *ΔvapC4* cells exhibited an increased tendency to form biofilm when subjected to a minimum of 30 mins of initial heat stress compared to WT and *ΔvapC4+vapC4* complementation strains **(Figure 8c)**. Similarly, a significant increase in biofilm formation was observed for *ΔvapBC4* cells when exposed to a minimum of 30 mins of initial heat stress **(Figure 8d)**. Moreover, a decreased tendency of biofilm production was seen in the heat-stressed Δ*vapBC4 strain* complemented with *vapC4.* All mutant strains and complemented strains utilized underwent PCR verification to confirm the presence or absence of the relevant genes (**Figure S4**). From these observations, we can speculate that heat stress-induced VapC4-mediated persister cells may not preferentially adopt a biofilm lifestyle as a coping strategy to heat stress, whereas heat stress-induced VapC4-deficient cells show an increased tendency to form biofilm as a coping mechanism to thermal stress.

**Figure 8.**
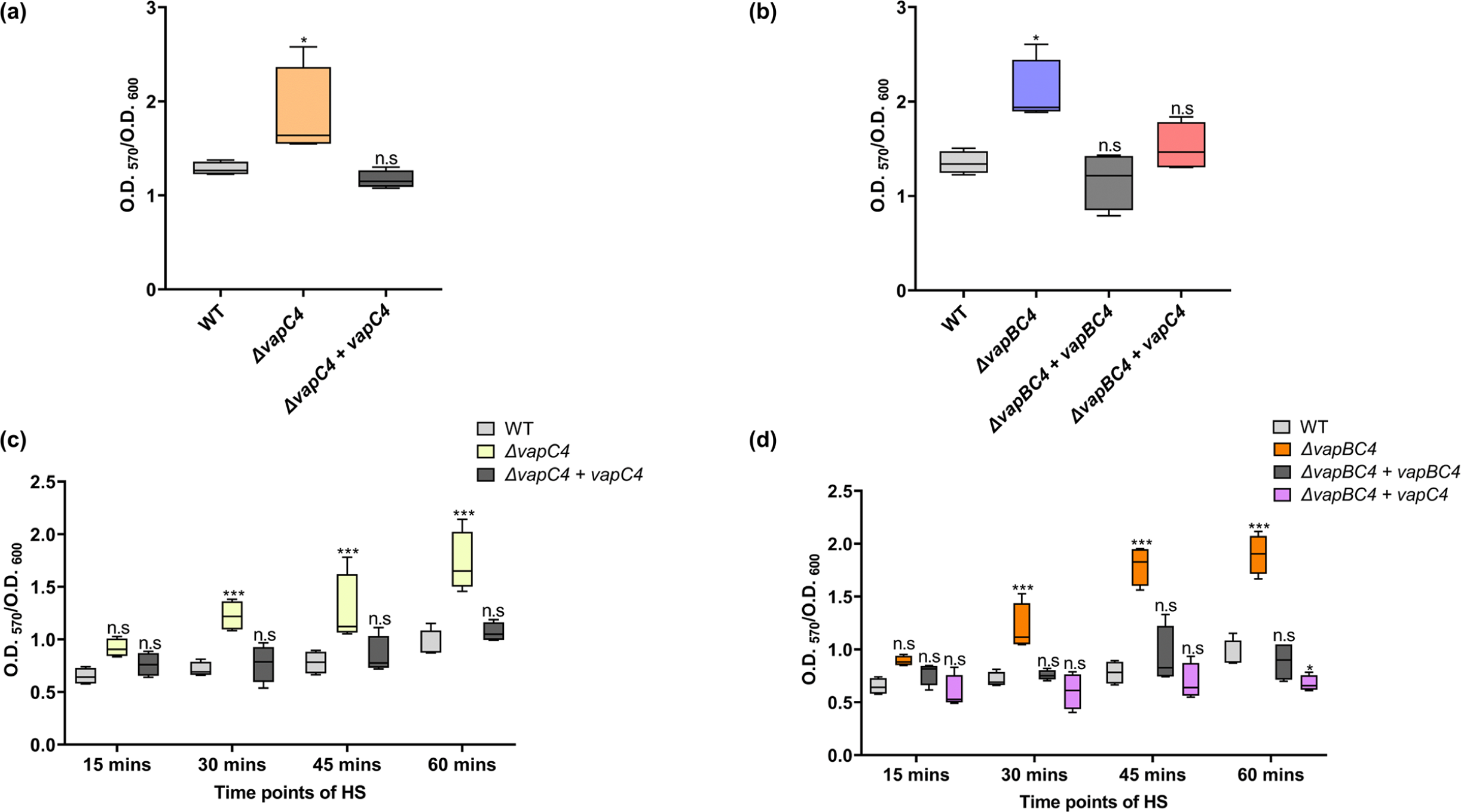
Impact of VapC4 toxin on biofilm formation. a) Box plot depicting normalised biofilm formation (OD_570_/OD_600_) for *ΔvapC4* cells vs. WT over two days. b) Box plot depicting normalised biofilm formation (OD_570_/OD_600_) for *ΔvapBC4* cells vs. WT and for *ΔvapBC4* + *vapC4* cells vs. WT over two days. c) Box plot showing normalised biofilm formation (OD_570_/OD_600_) for heat stress induced *ΔvapC4* cells vs. WT after varying time points of heat stress over 2 days. d) Box Plot showing normalised biofilm formation (OD_570_/OD_600_) for heat stress induced *ΔvapBC4* cells vs. WT and *ΔvapBC4 + vapC4 cells* vs. WT after varying time points of heat stress over 2 days. The statistical significance, P < 0.05, is indicated as *, P < 0.01 is indicated as **, and P < 0.001 is indicated as ***; n.s. denotes not significant.

## 4. DISCUSSION

The presence of a large number of *vapBC* toxin-antitoxin (TA) systems within the group of thermoacidocphilic archaea, including *S. acidocaldarius*, implies a possible role in adapting to their specific environments. Because bacterial TA systems are involved in programmed cell death or persister cell formation during stress conditions [2], we speculated that depending on the stress conditions, *S. acidocaldarius* might activate or deactivate specific *vapBC* systems to determine the fate of the cells. In our earlier research, we have demonstrated how the *vapBC* system plays a part in adapting to various stresses. We observed distinct sets of *vapBC* TA systems being upregulated in response to different stressors such as heat, oxidative conditions, and nutrient deprivation [15].

In this study, we showed that the heterologous expression of VapC4 in *E. coli* induced bacteriostasis, suggesting that the toxic influence of VapC4 extends beyond the archaeal domain. We demonstrated that VapC4 functions as a high-temperature catalyzed ribonuclease, validating the existence of a PIN domain fold. The ribonucleolytic activity of VapC4 might be involved in the inhibition of translation, which resulted in the repression of bacterial growth during its overexpression. Additionally, we have demonstrated that the VapC4 toxin hydrolyzes both rRNA and mRNA effectively *in vitro*. Further studies need to be performed to ascertain whether the hydrolysis by VapC4 in *S. acidocaldarius* is specific to the RNA sequence or structural folds. Also, subsequent in-vivo studies are required to ascertain whether ribosomes have any inhibitory role on rRNA cleavage by VapC4. The toxic activity of the VapC4 toxin is alleviated by its interaction with the cognate VapB4 antitoxin, resulting in the formation of a stable VapBC4 toxin-antitoxin complex-a crucial criterion for type II TA systems. This interaction might impede the access of the RNA substrate to the active site of VapC, as previously reported [16, 48]. Remarkably, we have also demonstrated the involvement of the vapC4 toxin in triggering persister-like cell formation in *S. acidocaldarius* during heat stress. This VapC4 toxin-induced susceptibility to heat stress at 85℃ in *S. acidocaldarius* can be attributed to the persister cell formation, where the cell enters a dormant state, allowing them to temporarily withstand and survive the challenging conditions posed by elevated temperatures, ensuring that the cells do not die. Persister cells in bacterial populations are characterized by their capacity to withstand challenging conditions, including heat stress, antibiotic exposure, and various environmental stressors [43–45]. These cells possess a unique capability to enter a dormant state, rendering them temporarily resistant or tolerant to external stresses that would generally eliminate actively growing cells [46]. The mechanisms underlying persister cell formation can involve various genetic, metabolic, and environmental factors contributing to their resilience in unfavorable conditions. Numerous studies have documented the induction of persister cell formation in bacteria through the TA system, enabling these cells to endure various stresses encountered in their natural environment [47–49]. These persister cells exhibit metabolic activity while inhibiting cell division, a strategic adaptation to cope with the imposed stress. The inhibition of translation by the VapC4 toxin through its ribonucleolytic activity during heat stress might be a cellular strategy aimed at conserving energy and resources while adapting to challenging conditions. Heat stress can lead to protein misfolding and denaturation, posing a significant threat to cellular functions [31, 49, 50]. By strategically inhibiting translation, the cell tries to minimize the production of new proteins, reducing the load on the cellular machinery. This reduction in protein synthesis serves several purposes. Firstly, it alleviates the burden on the cellular ribosomes, which may be compromised or overburdened under heat stress conditions. Secondly, limiting the production of new proteins helps prevent the accumulation of misfolded or damaged proteins, which could contribute to cellular dysfunction. Entering a persister state through translation inhibition might allow the cell to redirect its resources toward stress response mechanisms, such as chaperone proteins that aid in protein folding and repair [31, 49, 50]. This adaptive response helps the cell to better cope with the heat stress, promoting survival until more favorable conditions are restored. The proposed mechanism of persister cell formation by VapC4 toxin is illustrated as a model in (**Figure 9**). However, additional experiments are necessary to clarify the underlying physiology of this phenomenon. The persister cell formation has also been documented in *Haloferax volcanii* in response to starvation or exposure to lethal concentrations of different biocidal compounds [51]. On the contrary, VapC4 was identified as a negative regulator of biofilm formation, wherein the absence of the VapC4 toxin increased biofilm formation in *S. acidocaldarius*. Therefore, the free cellular VapC4 toxin during heat stress could cue the cell to opt for persister cell formation instead of biofilm formation. Persister cells exhibit temporary resistance to stressors without needing structural adaptations, allowing the cell to endure unfavorable conditions without forming complex structures like biofilms [52].

**Figure 9.**
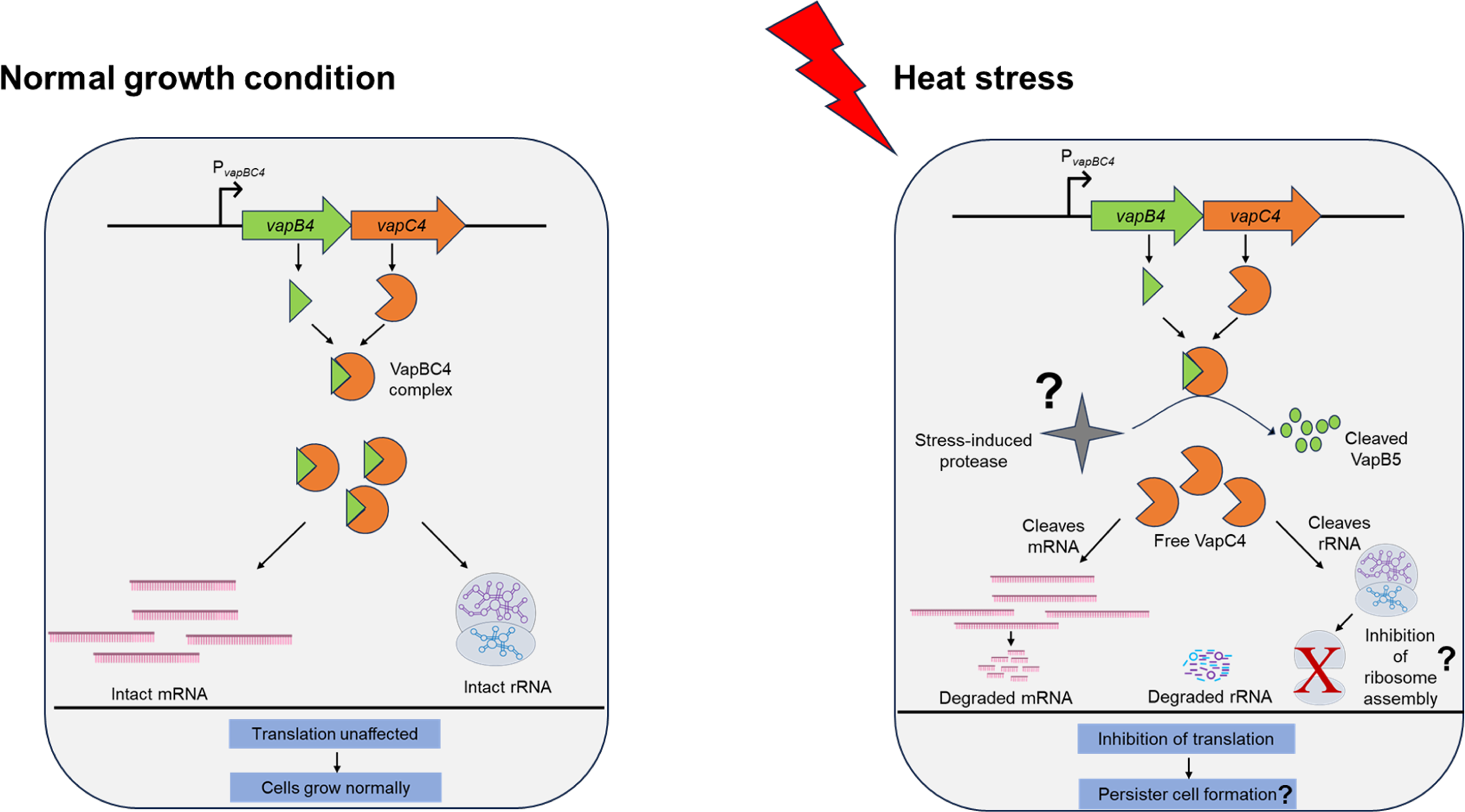
Proposed model showing the mode of action of the VapBC4 TA system during heat stress. Under the usual conditions, the ribonucleolytic action of the VapC4 toxin is blocked by the tight binding of VapB4 antitoxin, forming a stable VapBC4 TA complex allowing the cells to grow and divide normally. During heat stress, activated stress-induced protease (not yet known for archaea) might selectively cleave the VapB4 antitoxin. The unleashed VapC4 toxin, under stress conditions, can inhibit translation through two mechanisms: it can either directly hinder translation by cleaving mRNAs, or it can indirectly impede translation by cleaving 23S and 16S rRNAs, thereby destabilizing the interaction between the 50S and 30S ribosomal subunits (not yet known for archaea). Inhibition of translation serves as a vital mechanism for persister cell formation in response to stress conditions. This process is a strategic response to conserve energy and prevent the synthesis of potentially misfolded proteins under stress. The temporary halt in translation might redirect resources to stress response mechanisms, aiding in protein folding and repair. Persister cells, formed through this adaptation, enter a metabolically active but dormant state, allowing them to withstand challenging conditions until a more favorable environment is restored.

However, limitations exist within this study, including the lack of structural determination for VapC4 and the VapBC4 complex and the identification of the RNA cleavage site. In the future, it would be also interesting to examine whether VapC5 affects transcripts related to biofilm formation. Acknowledging these constraints, one could potentially unveil the underlying molecular mechanisms governing RNA substrate recognition and the binding dynamics with its corresponding antitoxin. Additional investigations are necessary to decipher the metabolic status and functionally characterize the persister cells of *S. acidocaldarius* to obtain a comprehensive understanding.

## Author contribution

A.G. and A.B. designed the study; A.B., A.R., C.B., J.D., and U.R.C. performed the experiments. A.G., A.B., A.R., and S.V.A. analyzed the data. A.G. and A.B. drafted the manuscript with inputs from all the authors. All the authors approved the submitted version.

## Declaration of competing interest

The authors declare that they have no conflict of interest with the contents of this title.

## Acknowledgment

The authors express gratitude for the intramural funding received from Bose Institute, India. AB acknowledges the fellowship support from CSIR (Council of Scientific and Industrial Research), Government of India, under File No. 09/015(0525)/2017-EMR-I. The authors extend their appreciation to EMBO (European Molecular Biology Organization) for granting AB the Scientific Exchange Grant (SEG 9915), enabling a 3-month visit to Prof. Dr. Sonja-Verena Albers’ laboratory at the University of Freiburg, Germany. ACR was supported by a Life Grant (AZ96727) by the VW Foundation awarded to SVA.

